# Generative enhancement of non-invasive datasets for motor brain-computer interface by synthesizing task-relevant neural signals

**DOI:** 10.1101/2025.10.12.681961

**Authors:** HongJune Kim, June Sic Kim

## Abstract

Despite the increasing adoption of deep neural networks (DNNs) in brain-computer interfaces (BCIs), developing high-degree-of-freedom (DOF) systems capable of decoding continuous movements, such as limb kinematics, remains a significant challenge. This difficulty stems from limited availability of task-specific neural features within individual neural signal datasets. To overcome this, we proposed a generative adversarial network (GAN) framework to enrich training features within neural signal datasets. Specifically, we synthesized artificial neural signal waveforms of the primary motor cortex (M1) from functionally related cortical regions, thereby enhancing neural datasets for improved motor kinematics decoding via DNN. Using magnetoencephalography (MEG) recordings during goal-directed arm-reaching tasks, our results showed that enhancing individual datasets with GAN-synthesized M1 signals significantly improved decoding performance by about 10% (*p* < 0.05). Such improved performance is sustained even in the absence of real M1 signals. We further generalized the proposed enhancement to the motor imagery BCI competition dataset to improve classification accuracy. Our results highlight the potential of signal-generative networks to improve and augment motor BCIs to achieve freely intended movements.

## I. INTRODUCTION

One of the core tasks within the brain-computer interface (BCI) could be decoding brain activities to predict behaviors. To decode brain activities more precisely, researchers have utilized recording systems with high spatial and temporal resolution, such as high-density electroencephalography (EEG) or magnetoencephalography (MEG), and employed machine learning (ML) frameworks. ML frameworks can computationally identify feature dimensions (or planes) within brain activities related to those behaviors [1]. Identifying appropriate features in conventional ML frameworks, such as support vector machine (SVM) or linear models, relies on an elaborate manual feature-engineering process. However, the feature-engineering process becomes challenging as data dimensions increase. Since brain activity comes from dozens to hundreds of channels with sampling frequencies of more than a thousand, it is sometimes challenging for conventional ML frameworks to identify appropriate feature dimensions in those multivariate signal data [2, 3].

To address this difficulty, researchers have developed strategies to elicit robust neural responses. For instance, subjects can perform tasks to induce P300 potentials or steady-state visually evoked responses (SSVEP) for several categories of intended behaviors. From those neural responses, ML frameworks can quickly identify significant features through manual feature-engineering processes and then classify several items or behaviors. Nevertheless, such strategies require devices and procedures to elicit responses (e.g., spontaneous gazing) [4]. In addition, degrees of freedom (DOF) are limited to less than a dozen categories [5, 6]. Thus, these strategies face challenges when BCIs need to decode high-DOF behaviors and carry out tasks continuously or in multiple ways, such as speech production, arm-reaching movement, or locomotion[5, 7]. Considering the characteristics of these behaviors, current strategies eliciting neural responses for several behavior categories could be inappropriate. Instead, an alternative approach that allows ML frameworks to decode high DOF movements from neural signals would be necessary.

The advancement of deep neural network (DNN) frameworks has recently contributed to the improvement of prediction accuracy in BCI [3]. DNNs are constructed by stacking multiple layers such as convolutional neural networks (CNNs), recurrent neural networks (RNNs), and dense layers by drawing inspiration from the hierarchical organization and learning mechanisms of biological neural systems [8]. By increasing the depth and width of networks, DNNs can learn higher-level and more abstract features that manual feature-engineering processes in conventional ML frameworks might struggle to extract. Thus, the current trend in BCI employs DNN frameworks to decode real-world data, such as classifying objects and time-series forecasting, replacing conventional ML frameworks.

Early approaches comprised DNNs with a few convolution layers and decoded robust EEG responses, such as P300, error-related negativity, sensory-motor rhythms, or SSVEP [9-11]. Results showed that the prediction accuracy improved compared to conventional ML approaches. Furthermore, since DNNs could directly decode spatio-temporal features within multi-channel neural data, task DOFs are significantly increased. For instance, RNN architectures [12, 13] could decode continuous movement kinematics and speech waveforms from neural signals [14, 15]. Moreover, large-scale DNNs comprising hundreds of layers and millions of parameters, such as Transformer [16], can significantly improve decoding performance compared to shallow architectures [17-19]. Meanwhile, reported decoding performance in BCI studies varies widely across recording modalities and task paradigms, with non-invasive EEG-based motor imagery typically achieving accuracies of 55–75% [2, 20-22], and higher performance can be achieved by increasing the modalities’ spatial and temporal resolution with MEG and ECoG [23, 24].

Nevertheless, DNN frameworks in BCI face two significant challenges. First, although deep and wide architectures allow us to decode complex neural signals directly and accurately, the number of training samples increases with the size of the architectures. However, acquiring neural data from individual subjects is resource-intensive and time-consuming [25]. Furthermore, augmenting datasets with data from multiple subjects may degrade BCI performance due to domain differences within- and across subjects [26, 27]. Thus, some researchers have transferred standard features derived from multi-subject data to train individual decoding models [20] or to augment datasets [25, 28]. However, the heterogeneous nature of neural signals across individual subjects and challenges associated with limited data acquisition continue to constrain the extent of performance improvements in BCIs. Second, although the prediction performance could be increased with signal-to-noise ratio (SNR), task-irrelevant components within signals could interrupt DNNs’ learning and cause a BCI-illiteracy issue [29, 30]. Due to such challenges, development in acquisition methods and employment of DNN frameworks might not directly improve BCI performances. Moreover, the neural signals, even within individuals, are inherently non-stationary and can vary over time and conditions due to factors such as fatigue, medication, distraction, and changes in cognitive or emotional state. These variations often degrade BCI’s performance, particularly under chronic application.

Most approaches have focused on increasing neural datasets by performing domain adaptation and data augmentation to address the first challenge. Domain adaptation methods can close distances between individual and target neural data domains [26, 27]. Meanwhile, data augmentation approaches have been tried to supply artificially synthesized neural data samples through generative neural networks. However, the effects of domain adaptation and data augmentation varied. Sometimes, the effects are negative [31], depending on the dataset and processing modes [32]. Besides, improvement rates did not linearly increase with the sample size of the synthesized dataset [31-33]. Such issues in data augmentation might be due to the second challenge: neural data could be accompanied by multi-scale artifacts such as non-neural noises and task-irrelevant cognitive states. Although the former could be filtered through conventional signal preprocessing pipelines, the latter were difficult to remove. Thus, those task-irrelevant features might mix in the data synthesized and muddle DNNs.

Enhancing the neural dataset with task-relevant neural features would effectively address this issue. Our previous work highlighted task-relevant neural features by decomposing DNNs within BCIs via an explainable AI (XAI) technique [21, 34]. Those identified cortical regions around motor-related areas contributed to decoding continuous hand movement kinematics. Although highlighting identified features contributed to the improvement of BCI performances [34], their reliability would be determined by the performance of DNNs. Furthermore, the task relevance could be varied across XAI methods [35]. Therefore, rather than highlighting features, supplying task-specific neural features directly into the dataset might be more effective.

Recent studies have increasingly explored generative modeling of neural signals, primarily focusing on data augmentation or reconstruction in EEG- and ECoG-based brain–computer interface (BCI) tasks [36]. These approaches have demonstrated that generative models can modulate classification robustness or signal quality by increasing training data diversity [25, 37-40]. However, most existing work treats synthetic neural signals as generic augmentations, without explicitly targeting task-specific cortical features or examining their contribution to movement decoding.

To do so, generative networks might help synthesize and supply task-specific features. They can learn the statistical representation of targeted data. Therefore, such networks can learn a mapping that translates source neural features into task-specific target representations [41]. Using the generative netowks’ characteristics, we thought to synthesize task-specific neural features, such as the motor cortex’s signals for motor BCI [34, 42].

Thus, we proposed a method that synthesizes task-specific neural features using a generative network from brain signal datasets to improve the decoding of hand-reaching movements toward four targets and motor imagery classes. We hypothesized that enhancing input datasets with task-specific neural signals of motor-BCIs synthesized by a generative adversarial network (GAN) could improve the decoding accuracy of a machine learning framework for predicting continuous hand-reaching kinematics.

## II. Dataset Information and Processing

### A. Magnetoencephalography datasets for motor-BCI

We employed MEG datasets from eight subjects for a motor BCI [35]. Each dataset included 306 channels of MEG (VectorView TM, Elekta Neuromag, Finland) and three-axis accelerometer signals recorded during arm-reaching movements toward four targets presented in 3D space. The experiment comprised two repeated sessions, including an inter-session breaktime. During each session, the subject repeatedly performed arm-reaching movements toward a target immediately after the onset of the visual cue. Those signals passed a 0.5 Hz high-pass filter. Non-neural signal components in MEG, such as oculomotor and electrocardiographic activity, were then removed using independent component analysis (ICA). MEG signals were then passed through band-pass filters from 0.8 Hz to 60 Hz. Likewise, accelerometer signals were passed through a band-pass filter from 0.5 Hz to 5 Hz. Filtered accelerometer signals were integrated twice to acquire directional hand movement positions over time. Lastly, both signals are resampled to 512Hz and sliced into epochs, -1 s to +2 s from the visual target onset (1536 time points per epoch). We obtained 240 trials (60 per the reaching direction) in the individual dataset, which were split into five folds for cross-validation.

Then, we estimated cortical source activity in the primary motor area (M1), medial intraparietal sulcus (mIPS), premotor area (BA6d), and visual motion area (MT) from 306-channel MEG via a dynamic statistical parametric mapping (dSPM) method [43]. These ROIs were participate in reaching movement production [42] (Supplementary Table I). To increase the signal-to-noise ratio (SNR) of estimation, we included a noise covariance matrix of MEG signals before visual target onset (-1s to 0s).

Since the dataset did not include individuals’ structural magnetic resonance (MR) images, we employed a FreeSurfer template MR [44-46]. Although the use of template MR is cost-effective [47], direct registration in the MEG space may cause misalignment. Therefore, we scaled the template MRI [48] to each individual’s digitized head shape to prevent misalignment.

### B. Electroencephalography (EEG) motor-imagery dataset

To validate the generalizability of the framework, we additionally employed a neural dataset recorded by different modalities and tasks. The BCI competition IV-2a dataset [49] includes 22-channel EEG signals from nine subjects during mental imagery tasks for four limb movement classes. We rejected the bad channels via visual inspection. The signals were passed through a 0.5Hz high-pass filter. Then, we corrected signal artifacts with ICA and passed them 1Hz to 30Hz band-pass filter. The signal was down-sampled to 128Hz and split into discontinuous epochs, -1s to 4s from visual cue onset (640 time points per epoch). We obtained 288 epochs (72 epochs × 4 classes) per individual subject. These epochs were split into four folds for cross-validation. In the analysis, we used five featured EEG channels in motor imagery classification [50]: Cz, Pz, Fz, C3, and C4.

## III. Model Description and Evaluation

### A. Generative adversarial network (GAN) for efficient and high-fidelity signal data synthesis

A generative adversarial network (GAN) is a robust architecture for synthesizing artificial data. The GAN comprises two mutually adversarial networks: the generator and the discriminator networks. The generator network synthesizes artificial (fake) data samples. In contrast, the discriminator networks learn to distinguish real from fake data samples in a given dataset. The generator network then tries to deceive the discriminator by learning the discriminator networks’ decisions and features [51]. Repeating such an adversarial learning procedure makes the generator produce more realistic fake data samples. Although previous studies have employed GANs to synthesize neural data samples [25, 28, 52-54], they have mainly utilized network architectures designed to generate image samples, even though the targeted data are neural signals.

Among the proposed architectures, a GAN architecture fulfilling the following two criteria was selected for high-fidelity neural signal synthesis. The first criterion was that the network should be specially designed to synthesize signal data, not images. The network architecture should consider the temporal dynamics of targeted signals represented in the waveform and spectral domains. Second, the GAN should comprise a fully convolutional architecture rather than an autoregressive one. Such a fully convolutional architecture has the advantage of decreasing computational cost during training and synthesis since the network can comprise fewer trainable parameters.

Therefore, we employed a HiFi-GAN architecture for neural signal synthesis. This neural vocoder considers spectral and temporal features in signal synthesis. Furthermore, it comprises a fully convolutional architecture suitable for real-time operation. The network showed promising performance in synthesizing artificial signals, such as speech and instrumental sounds, or in translating them into another characteristic. Likewise, we thought the HiFi-GAN would be used to translate neural signal characteristics.

### B. The network architecture of HiFi-GAN

Electrophysiological neural signals exhibit structures across multiple spectral and temporal scales, including low-frequency fluctuations to fast oscillatory components, which are associated with the behaviors. We employed a generative network that can capture the spectral and temporal dynamics of the neural signal.

The HiFi-GAN is a fully convolutional network that can synthesize signal spectrograms into raw waveforms [41]. This model includes three networks: (1) generator, (2) multi-period discriminator (MPD), and (3) multi-scale discriminator (MSD). The generator synthesizes fake signal waveform samples from the input spectrogram by integrating the product of multiple receptive fields (Fig. 1A). The discriminators identify real and fake waveform samples using long-term dependencies [55, 56] and periodic patterns within the signal samples [41] (Fig. 1B). The MSD enforces consistency of neural dynamics across long temporal contexts by evaluating signals at multiple resolutions, whereas the MPD explicitly emphasizes periodic patterns and fine-grained temporal structure [41].

**Fig. 1.**
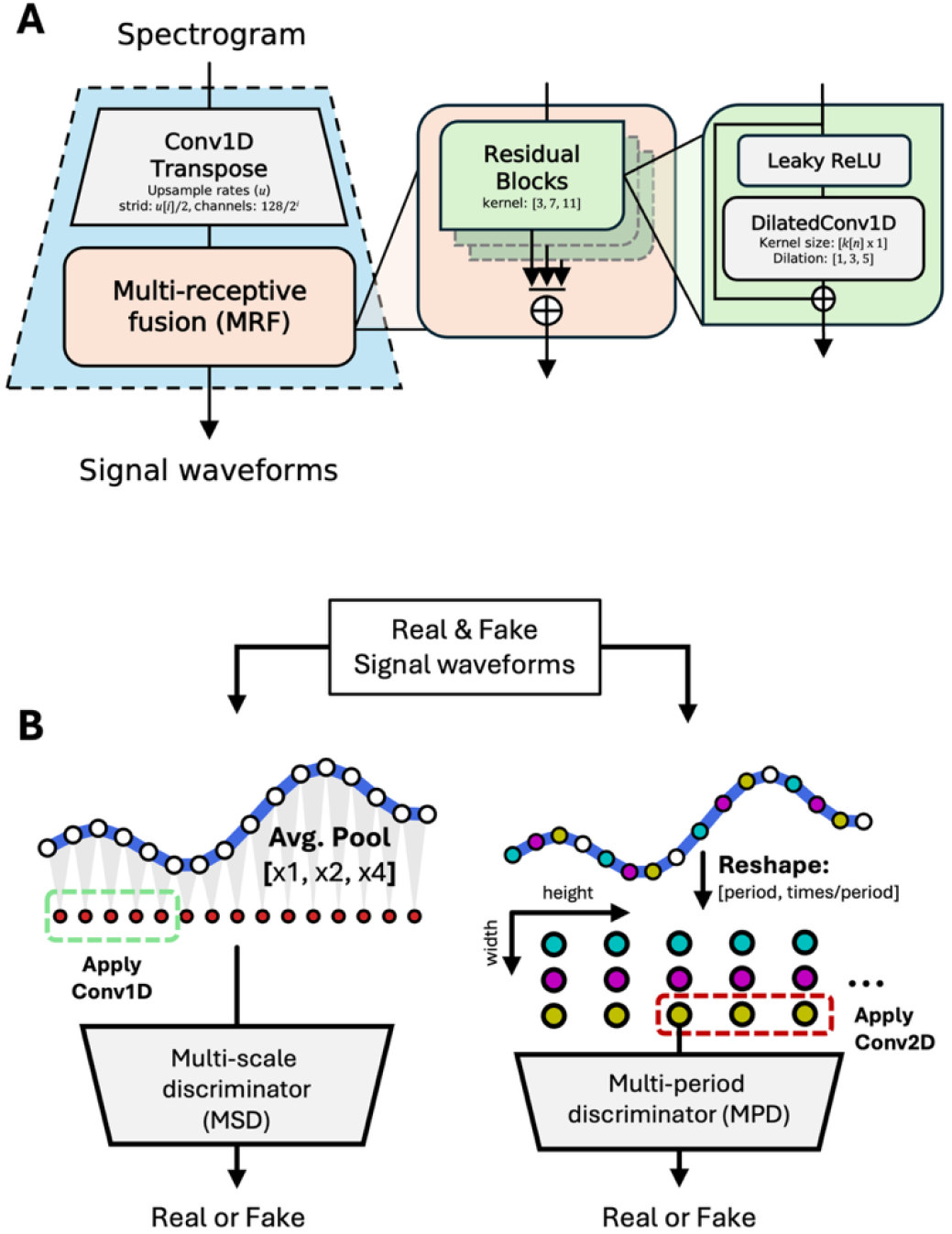
The HiFi-GAN architecture. (A) Generator network, (B) The discriminators: Multi-scale discriminator (MSD) (left) and multi-period discriminator (MPD) (right).

1. **Generator:** The signal spectrograms were upsampled through 1D transposed convolution (ConvTranspose1D). Following transposed convolution, the data passed through a multi-receptive-field fusion (MRF) module (Fig. 1A). MRF comprised multiple residual blocks with different kernel sizes and dilation rates and returned the sum of their outputs. Multiple kernel sizes and dilation rates can observe different temporal patterns in parallel, thereby forming diverse receptive fields. Furthermore, a dilated convolution design increased the receptive field size without increasing trainable parameters. The generator repeated the above process until the data shape matched the temporal resolution of raw waveforms.
2. **Multi-scale discriminator (MSD):** MSD comprised three sub-discriminator units for different signal scales. Each sub-discriminator received (1) raw signal waveforms, (2) x2 average-pooled smoothed waveforms, and (3) x4 average-pooled smoothed waveforms. Sub-discriminators processed those signals with strided and grouped convolutions and a leaky rectified linear unit (ReLU).
3. **Multi-period discriminator (MPD):** Sub-discriminator units within MPD reshaped raw waveforms into periodic signal blocks. When the period(p) was given, the sub-discriminator unit split the 1D raw waveform of t-length into a 2D periodic signal block of [t/p × p]. Then, 2D convolutions with a stack of strided convolution layers were applied. The convolution layers filtered the periodic signal block with [k × 1] kernels. Weight normalization was applied to the outputs of convolution layers.

We determined the HiFi-GAN’s hyperparameters by referring to the configuration of HiFi-GAN V2 [41]. In the generator, upsampling rates were [4, 4, 2, 2] with [8, 8, 4, 4] kernel sizes, scaled up to x64 from the spectrogram’s time samples until they reached raw waveforms. Residual block’s kernel sizes of each MRF module were [3, 7, 11]. Dilation rates of three convolution layers within the residual block were [1, 3, 5]. The total trainable parameters within the generator were 0.752 million.

Then, we downsized the number of filters and groups in the discriminators to reduce parameters, since the original version was too large to fit a single dataset. Thus, the sub-discriminator unit of MSD comprised 0.653M of trainable parameters. The first sub-discriminator for raw waveforms was applied spectral normalization [57], and others for smoothed waveforms were applied weight normalizations [58]. We comprised MPD with four sub-discriminator units of [2, 3, 5, 7]. Such an approach could prevent overfitting of discriminators and help identify oscillatory features within neural signals. Each sub-discriminator unit comprised 0.461M of trainable parameters.

### B. Optimization of the generative network

Firstly, the HiFi-GAN was trained using brain signals in the M1 as ground truth signals and the others as source signals. Ground truth signals were transformed into spectrograms using a short-time Fourier transform (STFT). The FFT size was 256, the window length was 256, and the hop size was 64. We then reduced the spectral dimension (frequency axis) using the 30 filter banks, which ranged from 1Hz to 60Hz. Thus, the input spectrogram’s tensor shape was [batch, 24-time points x 30 frequency channels].

The HiFi-GAN was optimized with the AdamW algorithm [59]. The learning rate was set to 0.0002. It then decayed by a rate of 0.999. We put the training iteration count to 20001 using two NVIDIA A100 GPUs. For the cross-validation, the dataset of individual subjects was split into five folds. We used 80% of the samples for training and 20% for testing and validation in each fold.

Likewise, we further evaluate the generalizability of the HiFi-GAN to the EEG motor imagery dataset. The model was trained using Cz signal. Since the EEG sampling frequency and resolution differed from those of MEG, we adjusted some parameters during training. We changed the upsamling rates to [2, 2, 2, 2], and kernel sizes to [4, 4, 4, 4]. The FFT size and window size were 64, and the hop size was 16. In addition, we down-sampled the spectral dimension to 20 filter banks, ranging from 1Hz to 30Hz. Lastly, the EEG dataset of individual subjects was validated with a four-fold cross-validation. Therefore, 75% of the samples were used for training, and 25% for testing.

### C. Neural signal synthesis evaluation

We tested the feasibility of neural signal waveform synthesis with HiFi-GAN. In the audio signal generation, the mean opinion score (MOS) was generally used as a metric for synthesis quality. It measures manual perceptual qualities [41, 56, 60] from the audience. However, such a metric could not be applied to neural signals. Thus, we evaluated generation quality by measuring representational similarity. First, we calculated the normalized cross-correlation (NCC) between the evoked (averaged) ground truth and synthesized neural signal waveforms. The metrics can intuitively show the similarities between evoked waveforms.

Nevertheless, the NCC could not assess similarity between distributions. Thus, the metrics should include the variances of those synthesized samples. We additionally calculated the mean L1-distance and the Fréchet distance between the ground truth and synthesized M1 signal samples. In addition, we included the Wasserstein-1 distance (Earth mover’s distance) between their power spectral densities (PSDs). Those metrics allowed us to compare the representative values and their variances.

The synthesis performance was compared against the baseline generative networks for EEG data: variational auto-encoder (VAE) [61], deep convolutional GAN (DCGAN) [54], Wasserstein GAN (WGAN-GP) [52], EEG-GAN [28]. Those referenced networks had been shown promising result on EEG data augmentation and neural vocoder. The EEG-GAN and VAE translated the input waveforms into targeted ones. The others synthesized the signal waveforms from the spectrogram.

### D. Decoding arm-reaching movement from the neural signals

We designed the RNN-based decoding model to predict hand-movement trajectories from neural signals. The model comprised a bidirectional long short-term memory (bLSTM) layer, a normalization layer, and two dense layers. The bLSTM layer processed sequential features within the time-series data. Outputs from the bLSTM layer were then normalized by the weight-normalization layer. Following the previous work [62], we preliminarily investigated the hidden unit sizes. We found that the hidden unit size of 100 showed the best performance on the sampled data. Then, the two dense layers regressed normalized sequential features to 3-axis coordination of hand-movement trajectories.

Before training DNNs, we processed neural and accelerometer signal data. Following previous studies [21, 62, 63], both real and synthesized neural signals were filtered with a 0.8 Hz to 8 Hz band-pass filter. We resampled those signals to 50 Hz for efficient computation. Then, neural signal samples were sequentially segmented into 200ms bins (10-time points per segment).

Likewise, we used five-fold cross-validation to test the entire decoding dataset. The neural signal and arm reaching movement dataset were split into five folds: 80% for training and 20% for testing. Total neural data segments per fold comprised a tensor of [26,880 segments, 10-time points, N-of feature channels] for training and [6,720, 10, N-of feature channels] for testing and validation. We decoded continuous three-axis arm reaching positions from neural signal segments. We removed linear drifts in trajectories using a linear detrending algorithm. Thus, the arm-reaching dataset comprised [26,880 segments x 3 axes] for training and [6720 segments x 3 axes] for testing and validation.

We fit the model with four dataset conditions: (1) baseline, (2) baseline with synthesized M1 from noises, (3) enhanced, and (4) enhanced without M1. The first baseline dataset only included actual neural signals of M1, BA6d, mIPS, and MT areas during hand movements. The second dataset contained all actual neural signals and one synthesized M1 signal from Gaussian noises. The enhanced dataset contained three synthesized M1 signals from mIPS, BA6d, and MT, along with actual neural signals. Lastly, we excluded actual M1 signals from the enhanced dataset to assess whether synthesized M1 signals could replace actual ones.

The objective function was the L1 distance between predicted and actual hand movement trajectories. The RNN-based model was optimized with the Adam optimizer with a 0.001 learning rate. We set training epochs to 100 with 32 mini-batch per iteration. The optimization ran on two parallel NVIDIA A100 GPU environments.

We further tested the proposed framework in a dataset with different modalities and cognitive tasks, and a machine learning model to assess generalizability. We decoded motor imagery classes from MI-EEG datasets with a support vector machine (SVM) classifier. Before fitting the SVM, we processed the EEG data through a band-pass filter of the alpha-band (7Hz to 12Hz) to extract motor-related features [2]. Like the MEG dataset, we fit the classifier with each dataset condition: (1) baseline, (2) enhanced. The baseline dataset included five EEG channels (Cz, Fz, Pz, C3 and C4). The enhanced dataset contained synthesized Cz signals along with EEG channels. We evaluated the four-class classification accuracy (chance level = 25%), F1-score, precision, and recall.

## IV. Results

### A. Neural signal synthesis performance of the HiFi-GAN

We evaluated the neural signal waveform synthesis performance of HiFi-GAN, compared to baseline models. We evaluated the signal synthesis performance of HiFi-GAN compared to the other baseline models. We synthesized M1 signal waveforms through each model from the mIPS signals. Table I shows the evaluation metrics between the ground truth and synthesized signal waveforms. Regardless of metrics, the HiFi-GAN outperformed the other baseline generative models. The synthesis performance improved by more than 15 times compared to the highest-performing baseline model, WGAN.

**Table 1.**
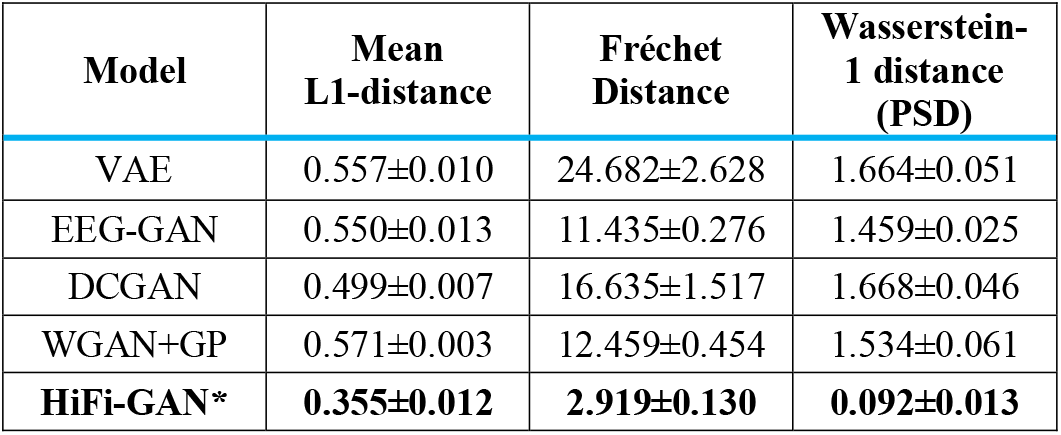
Comparing Generative Synthesis Performance to the Baseline Models.

### B. Representational similarity between ground truth and synthesized neural signal waveforms

Then, we assess the representation of neural signal waveforms synthesized through HiFi-GAN. Fig. 2. illustrates subject 1’s evoked signal waveforms and standard deviations of M1 (*x*, ground truth), mIPS (*s*, source), and synthesized M1 from the source (*G(s)*) and Gaussian noises (*G(cp)*). Results showed that the statistical representation of the synthesized M1 signal followed the ground truth (Fig. 2A). Furthermore, we identified the difference depending on the arm-reaching direction in evoked signals of the M1 and the source. Such differences were also identified in the evoked synthesized M1 signals. They followed ground truth ones (Fig. 2B). In contrast, differences in the direction of evoked signals were not observed in synthesized signals from Gaussian noise.

**Fig. 2.**
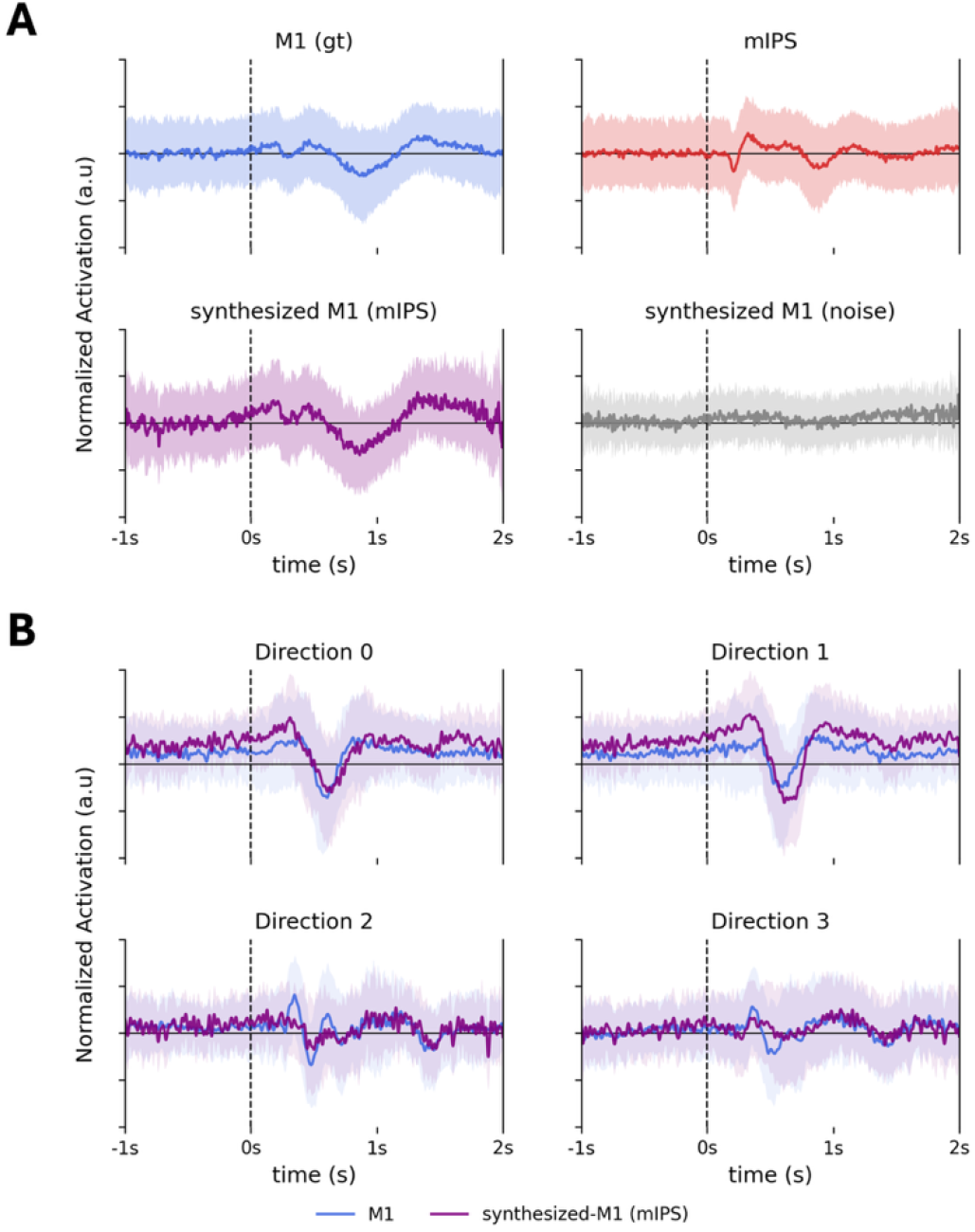
Signal waveforms of M1 (ground-truth), mIPS (source), and synthesized M1 signals from source and Gaussian noises. (A) Evoked signal waveforms and distribution of subject 1. Waveforms of all subjects are presented in Supplementary Figure 1. (B) Evoked signal waveforms and distributions of signals per hand-reaching direction of subject 5.

Quantitative results revealed that the NCC between those evoked signal waveforms showed a strong positive correlation between the synthesized M1 and ground truth signal waveforms under all conditions (Fig. 3). NCC results between ground truth and synthesized M1 signal waveforms from BA6d and MT are shown in Supplementary Figure 2. In every subject, the NCC peaked at 0s-lag, and the overall NCC waveform followed autocorrelation (NCC(*x*, G(*s*))_*t*=0_ > .716). Along with NCC, the FD between all those signal pairs showed that the normalized FD was closer between ground truth and generated signals than between M1 and source signals (Fig. 3). In addition, the FDs between synthesized and source signals (BA6d, mIPS, and MT) showed considerable distances. Normalized FDs between signal pairs in every subject are shown in Supplementary Figure 3.

**Fig. 3.**
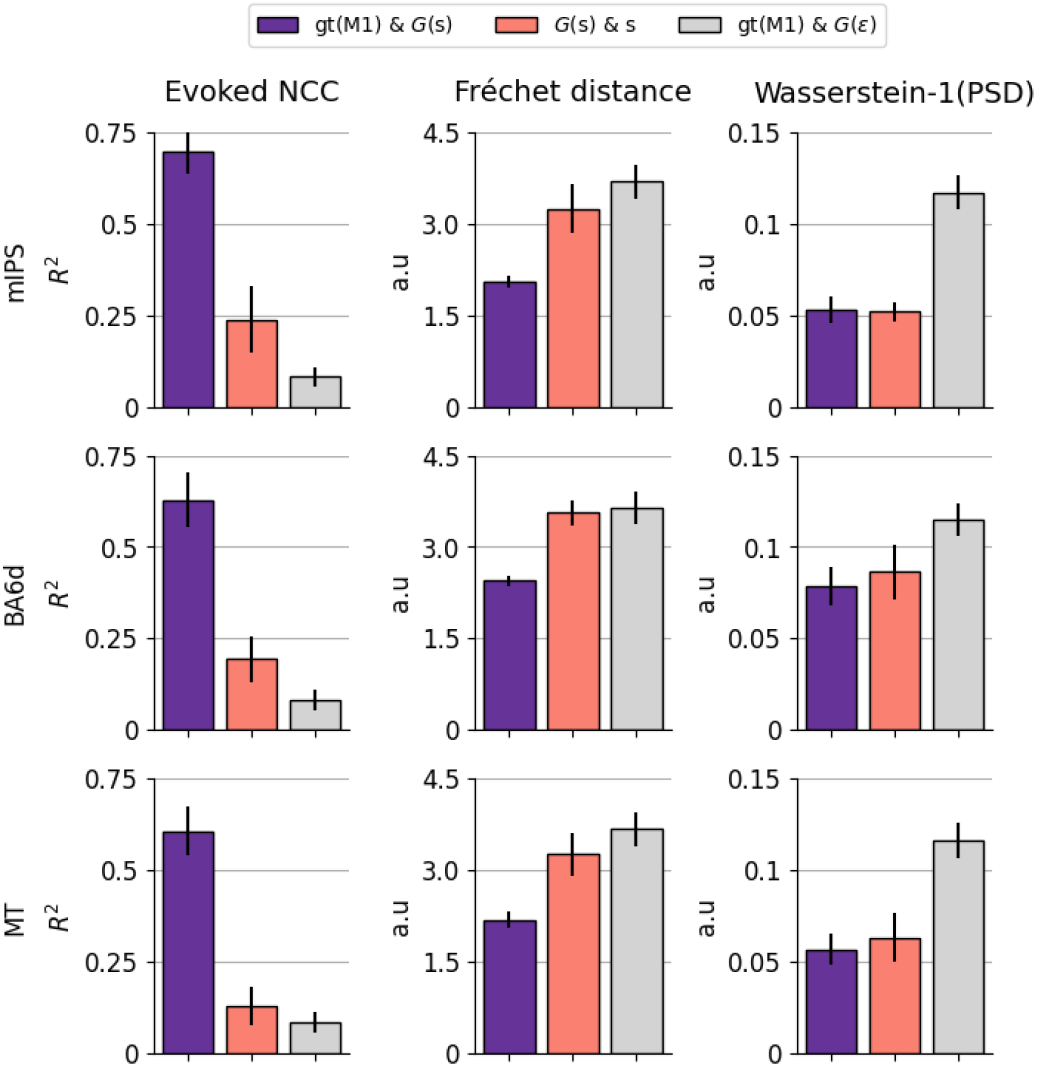
Quantitative metrics between the ground truth (M1), sources (mIPS, BA6d, and MT), and synthesized M1 signals using sources and Gaussian noises (*ε*).

Along with the signal waveform domain, we also tested the frequency domain representations of those synthesized signals. To access the frequency domain, we computed PSDs of the ground truth and the synthesized M1 signals using the sources and Gaussian noises. Then, we compared the Wasserstine-1 distances between them. The result shows that the PSD of the synthesized M1 from source signals matched ground truth; in contrast, the Gaussian noise did not (Fig. 4). Quantitatively, the Wasserstein-1 distance to the ground-truth signal was significantly closer in synthesized M1 using the source (0.053 ± 0.007) than using the noise (0.117 ± 0.009) (*t*(7) = -3.902, p < .01, 95% CI = [-0.102 0.025]) (Fig. 3).

**Fig. 4.**
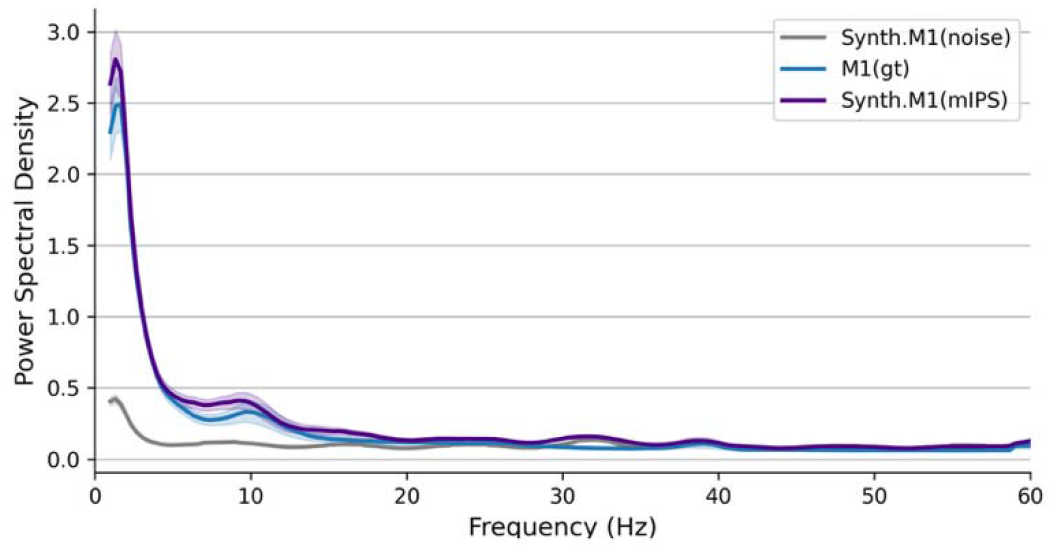
Averaged Power Spectral Density (PSD) of the M1 (ground truth) and synthesized M1 using source (mIPS) and using Gaussian noise.

### C. Decoding of arm-reaching trajectories

Regarding decoding performance, the enhanced dataset showed the highest accuracy (*r* = 0.401 ± 0.0207), followed by baseline (*r* = 0.366 ± 0.0201), enhanced dataset without actual M1 (*r* = 0.362 ± 0.025), and baseline with synthesized M1 from noises (*r* = 0.263 ± 0.019) datasets (Fig. 5A). We tested differences between those conditions with ANOVA. Results demonstrated significant differences between datasets (*F* (3, 21) = 25.049, *p* < 0.001, *η*^2^=0.38). Pair-wise t-tests were then performed for post hoc analysis. All p-values of t-tests were corrected via Benjamin-Hochberg false discovery rate (FDR) correction (Table II).

**Table 2.** Multiple Comparison Between the Dataset Conditions for Arm-reaching Prediction.

| Datasets | | A-B<br>(Effect<br>size $r$) | 95% CI | $t$-values<br>($df = 7$) |
| --- | --- | --- | --- | --- |
| A | B |  |  |  |
| Enhanced | Enhanced w/o M1 | 0.172<br>(0.63) | [0.10, 0.25] | 5.45*** |
| Baseline + Synth. M1 (noise) | Enhanced w/o M1 | -0.468<br>(0.91) | [-0.71, -0.23] | -4.60*** |
| Baseline | Enhanced w/o M1 | 0.023<br>(0.08) | [-0.14, 0.18] | 0.34 |
| Baseline + Synth. M1 (noise) | Enhanced | -0.640<br>(0.96) | [-0.85, -0.43] | -7.10*** |
| Baseline | Enhanced | -0.149<br>(0.64) | [-0.28, -0.02] | -2.66** |
| Baseline | Baseline + Synth. M1 (noise) | 0.491<br>(0.93) | [0.27, 0.72] | 5.15*** |

\* $p < .05$ , \*\* $p < .01$ , and \*\*\* $p < .001$ (FDR corrected)

**Fig. 5.**
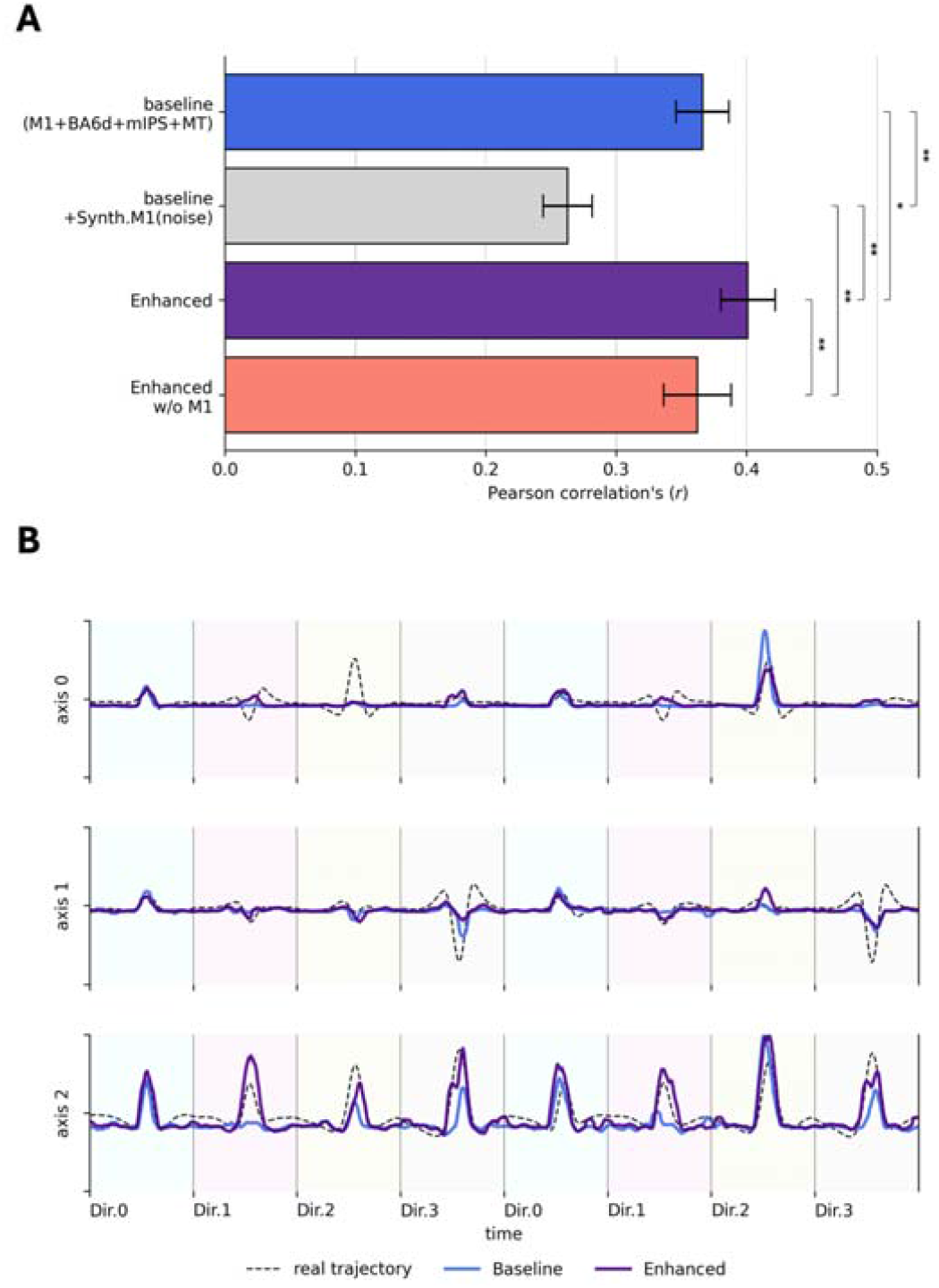
Decoding performances of neural signal dataset conditions. (A) Pearson’s correlation (r) between actual and predicted continuous hand movement trajectories (randomly sampled in each direction). Asterisks show statistical significance of difference between two dataset conditions. *, p < 0.05; **, p < 0.01; ***, p < 0.001 (Hochberg-Benjamin FDR corrected). The subject-wise performance is provided in the Supplementary Figure 4. (B) time course trajectories of actual (dotted) and predicted (solid lines) trajectories

Time-series DNNs trained on the enhanced dataset significantly outperformed those trained on other datasets. In contrast, the baseline dataset, appended with synthesized M1 signals from Gaussian noise, significantly underperformed. The baseline and enhanced datasets without actual M1 signals showed similar decoding performances. Their difference was insignificant. Representatively, decoded hand-reaching trajectories from the enhanced dataset closely matched those from the actual dataset (Fig. 5B).

### D. Generalizability to the other dataset

Lastly, we tested whether the proposed framework could be generalized to another dataset acquired through a different modality and for another task. First, we synthesized the neural signals of the artificial Cz channel through HiFi-GAN using the other EEG channels (Pz, Fz, C3, and C4). In line with the MEG result, the synthesized EEG waveforms of Cz, followed the ground-truth waveforms (Fig. 6A). We further measured the distances metrics between them. The FD results showed that the synthesized signal waveforms were significantly closer to the ground truth than the source signals (*t*(8) = 7.46, *p* < 0.01, 95% CI = [0.43 0.82]). Likewise, the Wasserstein-1 distances between their PSDs yielded the same result (*t*(8) = 2.62, *p* < .05, 95%CI = [0.01, 0.04]) (Fig. 6B). Furthermore, the 0s-lag NCC between the ground truth and synthesized Cz (NCC(*x*, G(*s*))_*t*=0_ > .736) was higher than between the synthesized and the source (NCC(*s*, G(*s*))_*t*=0_ < .132).

**Fig. 6.**
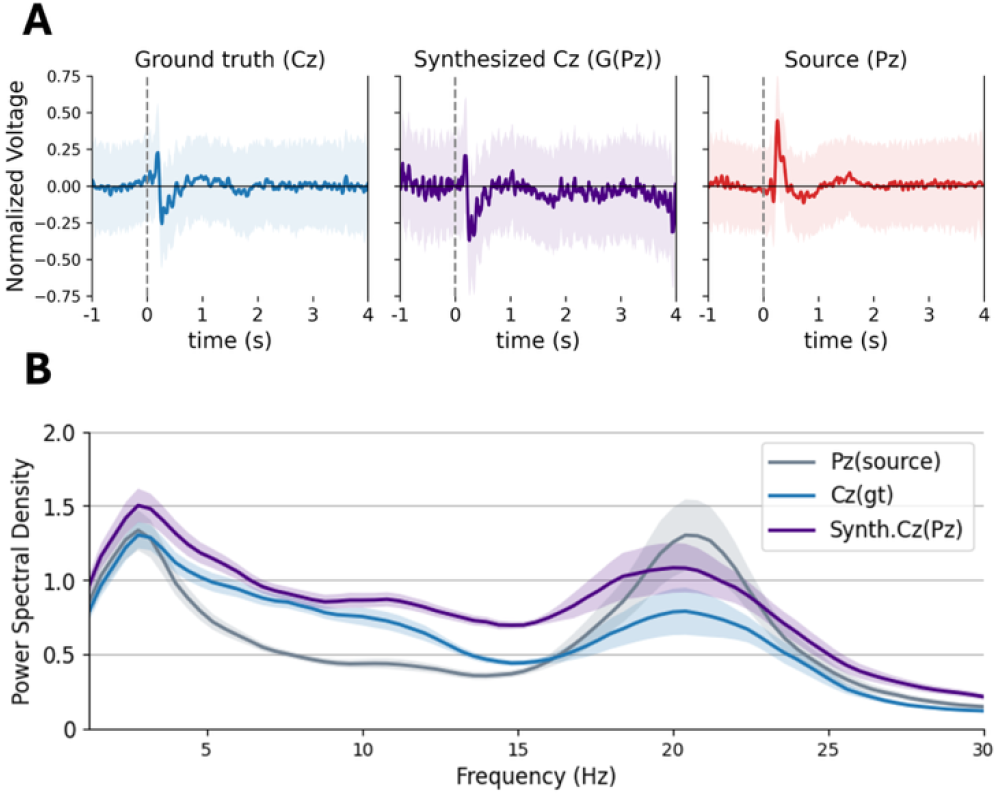
Generative synthesis of EEG signals during the motor imagery tasks. (A) Evoked EEG signals and their distributions in Cz (ground truth), Pz (source), and synthesized Cz channels using source. (B) Mean power spectral density (PSD) and their distributions of them

Then we tested the decoding performance using the synthesized Cz signals. We classified the four classes of motor imagery content from each dataset using a support vector machine (SVM) classifier. We tested the decoding performance by a four-fold cross-validation. The mean decoding performance with the enhanced dataset was 60.72% ± 4.19%. It was significantly higher than the baseline dataset, 58.91% ± 4.66% (*t*(8)=2.33, *p* < .05, CI=[0.73, 3.75]). Such a result could be found in the other metrics, precision (*t*(8) = 2.62, *p* < .05, 95%CI = [0.01, 0.04]), F1-score (*t*(8) = 2.12, *p* < .05, 95%CI = [0.01, 0.05]), and recall (*t*(8) = 2.39, *p* < .05, 95%CI = [0.01, 0.04]) (Supplementary Table II).

## V. Discussion and Conclusion

One of the main objectives of motor-BCI is to decode neural signals and predict time-series movement using DNNs. The present study aimed to facilitate DNN training with individual datasets by enhancing their feature dimensions using task-specific neural data samples. Since the task was to achieve center-out hand-reaching mov. ements, task-specific neural signals should directly represent signal characteristics within the motor cortex. While macroscopic signals from the primary motor cortex are inherently limited in precisely determining movement direction [34], effectively transforming and incorporating directional information from other brain areas to expand feature space could substantially improve movement prediction accuracy. Thus, we employed a GAN designed to learn spectral and temporal dynamics within the M1’s signal waveforms. Through the HiFi-GAN, artificially synthesized M1 signal waveforms were generated with those from secondary motor-related areas such as PRR, BA6d, and MT. Results showed two significant findings. First, the GAN could synthesize task-specific neural signal waveform samples from other task-related sources based on their spectral and temporal dynamics. Second, enhancing the neural dataset by supplying synthesized neural signal waveforms within feature dimensions could facilitate DNN training and improve its performance.

These findings indicate that the generative model does more than replicate target signals. It also actively reveals task-relevant features embedded within the neural data. Because macroscopic brain signals result from mixed activations from a vast number of neurons, each channel or source space inherently contains a wealth of intermingled information. The generative model can effectively extract task-related signals from such complex data. This finding is consistent with observations in generative applications for speech and image synthesis, where fusion of original and target signal characteristics gives rise to new information [55, 64]. Consequently, our results suggest the potential of using generative AI not only to replicate neural signals but also to deepen our understanding of underlying neural mechanisms and advance neural engineering applications.

### A. A generative network framework to synthesize high-fidelity neural waveforms for a BCI system

Previous approaches have adopted generative model architectures validated on image dataset synthesis, such as Wasserstein GAN [28, 52], variational autoencoder [65], and StarGAN [25]. However, their data types are temporal neural signals. Those approaches are practical for synthesizing the static ‘imaging’ aspect of neural signals, such as topological representations for SSVEP or P300, to classify discrete targets. However, such approaches could not synthesize neural signal waveforms. Unlike static image data, signal waveforms can be characterized by spectral and temporal dynamics in the time-frequency domain.

The HiFi-GAN architecture considers both spectral and temporal features. Thus, it can synthesize samples reconstructing signal waveforms. The training of the generator and discriminator networks includes loss between waveforms and the time-frequency domain, representing spectral and temporal dynamics. Furthermore, losses in MSD units preserve temporal characteristics, even in the low-resolution representation. In contrast to previous signal synthesis GAN [60], which uses only MSD, MPD units also consider features present in periodic signal patterns during discrimination. Loss terms with the MPD’s decision and features help the generator synthesize neural signals more realistically. As a result, the HiFi-GAN outperforms other baseline models, such as EEG-GAN, by more than X10 in neural signal synthesis.

Moreover, since the network comprises fully convolutional architectures, the computational cost of training and synthesis is lower than that of autoregressive models. The trainable parameters in the architecture are small enough to train on individual datasets (less than 0.7M). Since we could train RNN models comprising more than three million parameters on the same MEG dataset in previous studies [20, 21, 34], such networks within the GAN can also be trained with individual datasets. Moreover, synthesis time per trial requires only a few milliseconds in the NVIDIA GPU environment. We speculate that such a framework can be implemented in a real-time BCI system.

To synthesize motor neural signals, we used cortical source signals estimated in secondary motor-related areas, such as BA6d, MIP, and MT. The synthesized M1 signal waveforms were compared with those signals, followed by a statistical representation of ground truth signals, whereas Gaussian noise signals were not. Furthermore, the representational similarity in evoked waveforms and their distribution is preserved in each direction. Evoked waveforms of synthesized M1 signals for each direction also followed statistical representations of ground truth (Fig. 2B). In contrast, synthesized signals with Gaussian noise did not show differences in each direction (Fig. 2B). Moreover, the frequency domain representation also shows that the PSD of the synthesized M1 signals using the secondary motor area matched the ground truth’s features for motor controls, such as μ-band oscillation (7Hz - 11Hz) [23, 63, 66].

Regarding the findings of this study, those signals in secondary motor-related areas could provide conditional information to the GAN. Such speculation can be supported by the brain mechanism involved in the production of hand-reaching movement. The mIPS is called the parietal reaching region (PRR) within the posterior parietal cortex. It encodes both pointing targets (goals) and preparation for reaching movement [42]. In addition, functional modulation of the area can affect the success of the reaching movement [67]. Moreover, the BA6d within the premotor area receives neural codes for reaching movement from mIPS through the parieto-frontal network [68, 69]. It encodes discrete components for the upcoming movement rather than a continuous one [70]. The MT area shows a motion-dependent representation [71] and provides manual following responses, crucial for controlling body position during reaching [72]. Thus, neural codes for directional movements embedded within source signals offer conditional information to the GAN, allowing the network to synthesize M1 signal waveforms based on that information. The method also has advantages in computation and application, since no conditional codes are required in the generative network for synthesis. It just needs neural signals in functionally related areas.

### B. Improving high-DOF motor-BCIs with an enhanced neural signal dataset with task-related synthesized neural signals

This study aims to facilitate DNN training with a neural dataset enhanced with synthesized task-specific neural signals. As proof, enhancing the neural signal dataset with synthesized M1 signals from cortical sources should improve time-series DNN’s decoding performances. Results showed that decoding performance improved by 9.45% when the DNN was trained on the enhanced dataset compared to the baseline. This improvement validates that synthesized task-specific neural data, such as M1’s neural activity for continuous movements [21, 70], can help DNN identify more accurate hidden spatial and temporal features in training.

We speculate that synthesized M1 signals from motor-related cortical sources can help the DNN learn hidden features embedded within the M1 for continuous movements. Those synthesized M1 signals are not mere copies or reproductions of the ground truth. The generator network synthesizes unseen M1 signal waveforms with the source signals following a statistical representation of the ground truth. Such synthesized M1 signal waveform samples may supply neural codes for continuous movements embedded within M1 and facilitate learning. Nevertheless, actual M1 signals could still significantly affect DNN training since excluding the area reduced decoding performances.

Meanwhile, synthesized M1 signals from secondary motor-related areas can substitute for M1’s functional role. Although excluding actual M1 signals from the enhanced dataset reduced decoding performance, it remained comparable to that of the baseline dataset. There was no significant difference between them. This result suggests that the generative network can complement neural activity from functionally related brain areas. For instance, intracranial electrodes such as intracranial EEG (iEEG) and microelectrodes can provide high spatial and temporal resolution neural signals for invasive BCI systems [73]. However, intracranial electrodes can only measure focal brain regions [74, 75]. Thus, capturing all task-related regions for running BCI systems with modalities would be challenging. By using our framework, we speculate that the generative network may help reconstruct neural signals within task-specific areas that intracranial electrodes cannot measure, thereby facilitating the development of invasive BCI systems. The applicability to other modalities was tested by synthesizing EEG signals in the public dataset.

An important direction for future investigation is to examine how decoding performance varies with the proportion of real and synthesized neural data. While the present study demonstrates that generative synthesis can effectively enhance real neural signal datasets, systematic analysis of substitution ratios (e.g., partial replacement of real data with synthesized data) may further clarify the limits and applicability, particularly in low-data or longitudinal BCI settings.

### C. Generalizability to the other neural signal modality and the BCI tasks

Lastly, we examined whether the proposed generative enhancement framework remains effective under another dataset condition. We synthesized EEG signal waveforms of the Cz channel using the other channels during the motor imagery task. Then, imagined movements are classified from EEGs using a machine-learning model. In line with the MEG arm-reaching dataset, the generative model was able to synthesize signal waveforms under the EEG and the motor imagery conditions. In addition, appending the artificially synthesized Cz signals improved the classification accuracy. These results suggest that the generative synthesis framework may capture task-relevant neural features associated with both imagined and executed movements. These results provide preliminary evidence that the proposed framework can be applied across different recording modalities and task contexts within the evaluated conditions.

### D. Limitations

Our neural signal synthesis and dataset enhancement framework was only tested on task-specific data, including noninvasive neural signals and arm-reaching movement datasets. Therefore, it may be hard to guarantee that our framework can be generalized to other BCI environments. Notably, since decoding performance with synthesized neural signals from random noises is too poor to be utilized for training the BCI system, signals from task-relevant areas should be used in synthesis. Further research should ensure that the synthesis and enhancement framework is practical for other tasks, such as decoding speech or motor imagery.

Although we showed that the proposed framework may be robust to inter-session variability, it was only tested in the two consecutive experimental sessions conducted on the same day. Therefore, the result can be limited to addressing chronic decoding caused by drifts in neural states over days of long-term use and by fatigue, medication, or changes in cognitive states. While the proposed framework did not directly address this challenge, it may serve as a foundation for future adaptive approaches, such as fine-tuning strategies, to improve robustness over long-term application.

Although enhancing the EEG dataset with synthesized Cz signals improves classification performance, this improvement is less than in the MEG arm-reaching dataset. The EEG has a lower SNR than MEG [73, 76]; thus, it contains more task-irrelevant components in the signals. Although we rejected and corrected the non-neural noise components using digital signal filtering and ICA, these components might still be mixed into the neural signal synthesis procedure.

### D. Conclusion and future perspectives

This study demonstrates that enhancing the neural dataset’s feature dimensions with synthesized task-specific neural signal waveform samples can facilitate DNN training and improve decoding performance. Results of this study suggest that a signal synthesis GAN can synthesize M1 signal waveform samples based on neural codes for reaching movement embedded within other motor-related source signals. Appending synthesized M1 signals can provide abundant features for continuous movement in neural datasets and enable DNNs to decode arm-reaching movement trajectories toward targets. Generative network frameworks that can enhance neural datasets will effectively complement the current challenge in DNN-based BCI systems, which are trained on small individual datasets. In addition, the neural signal enhancement can be effective in the motor imagery task and the EEG dataset. Thus, the frameworks may be generalized to other neural signal modalities and task conditions.

