## Supplementary figures for "Generative enhancement of non-invasive datasets for motor brain-computer interface by synthesizing task-relevant neural signals"

SUPPLEMENTARY TABLE I.

DESCRIPTIVE INFORMATION ABOUT REGIONS OF INTEREST (ROIS)

| **No.** | **Regions of interest (abbreviation)** | **MNI coordination (mm)** | | | **HCPMMP1 parcellation labels** | **Descriptions** |
|---|---|---|---|---|---|---|
| | | ***x*** | ***y*** | ***z*** | | |
| 1 | Primary motor cortex (M1) | -35.28 | -19.43 | 65.04 | L_4_ROI-lh | Primary motor area (Brodmann area 4, BA4) |
| 2 | Brodmann area 6 dorsal (BA6d) | -33.75 | -13.41 | 63.69 | L_6d_ROI-lh | Premotor area |
| 3 | Medial Intraparietal sulcus (mIPS) | -20.41 | -61.56 | 42.15 | L_MIP_ROI-lh | Parietal reaching region (PRR) |
| 4 | Middle Temporal area (MT) | -45.16 | -72.61 | 6.42 | L_MT_ROI-lh | Visual motion area |

SUPPLEMENTARY TABLE II.

CLASSIFICATION ACCURACY ON EEG-DATASET

| Subject | Features | accuracy (%) | Precision | F1-score | Recall |
|---|---|---|---|---|---|
| A01 | **Enhanced** | **64.58%** | **0.63** | **0.67** | 0.65 |
| | Baseline | 62.80% | 0.61 | 0.65 | 0.63 |
| A02 | **Enhanced** | **62.85%** | **0.63** | **0.63** | 0.63 |
| | Baseline | 60.42% | 0.60 | 0.61 | 0.6 |
| A03 | **Enhanced** | **61.46%** | **0.60** | **0.61** | 0.61 |
| | Baseline | 61.11% | **0.60** | 0.6 | 0.61 |
| A04 | **Enhanced** | **70.83%** | **0.70** | **0.71** | 0.71 |
| | Baseline | 69.44% | 0.69 | 0.70 | 0.69 |
| A05 | **Enhanced** | **39.24%** | **0.38** | **0.38** | 0.39 |
| | Baseline | 32.29% | 0.30 | 0.31 | 0.32 |
| A06 | **Enhanced** | **42.01%** | **0.41** | **0.43** | 0.42 |
| | Baseline | 40.62% | 0.40 | 0.42 | 0.41 |
| A07 | Enhanced | 61.11% | 0.60 | 0.61 | 0.61 |
| | **Baseline** | **62.5%** | **0.62** | **0.62** | 0.62 |
| A08 | **Enhanced** | **66.32%** | **0.66** | **0.67** | 0.66 |
| | Baseline | 63.19% | 0.62 | 0.63 | 0.63 |
| A09 | **Enhanced** | **78.12%** | **0.78** | **0.79** | 0.78 |
| | Baseline | 77.78% | 0.77 | 0.78 | 0.78 |
| Total | **Enhanced** | **60.72%±4.19%** | **0.611±0.043** | **0.599±0.042** | **0.607±0.042** |
| | Baseline | 58.91%±4.66% | 0.591±0.044 | 0.578±0.048 | 0.589±0.046 |

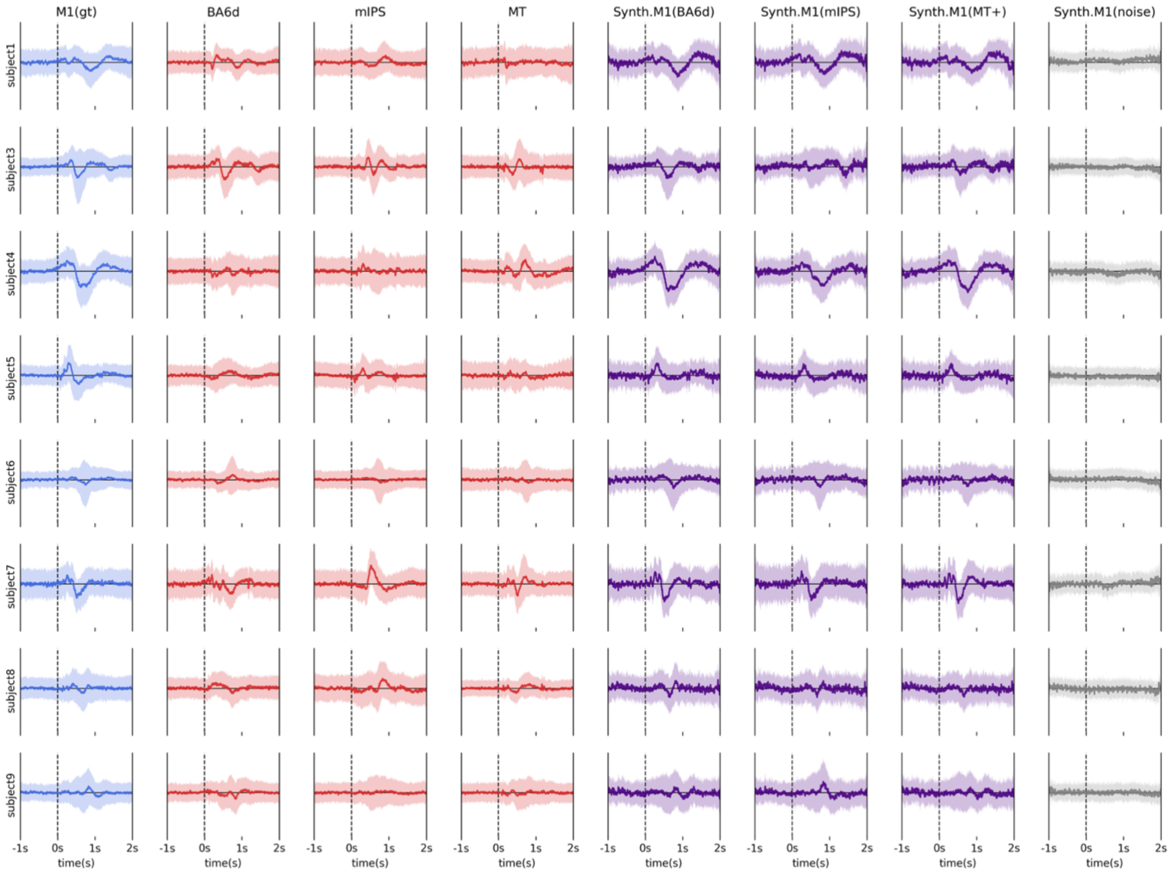


**Supplementary Figure 1.** The subject-wise evoked responses and distributions of all neural signals (actual and synthesized) in the arm-reaching dataset (BCI).

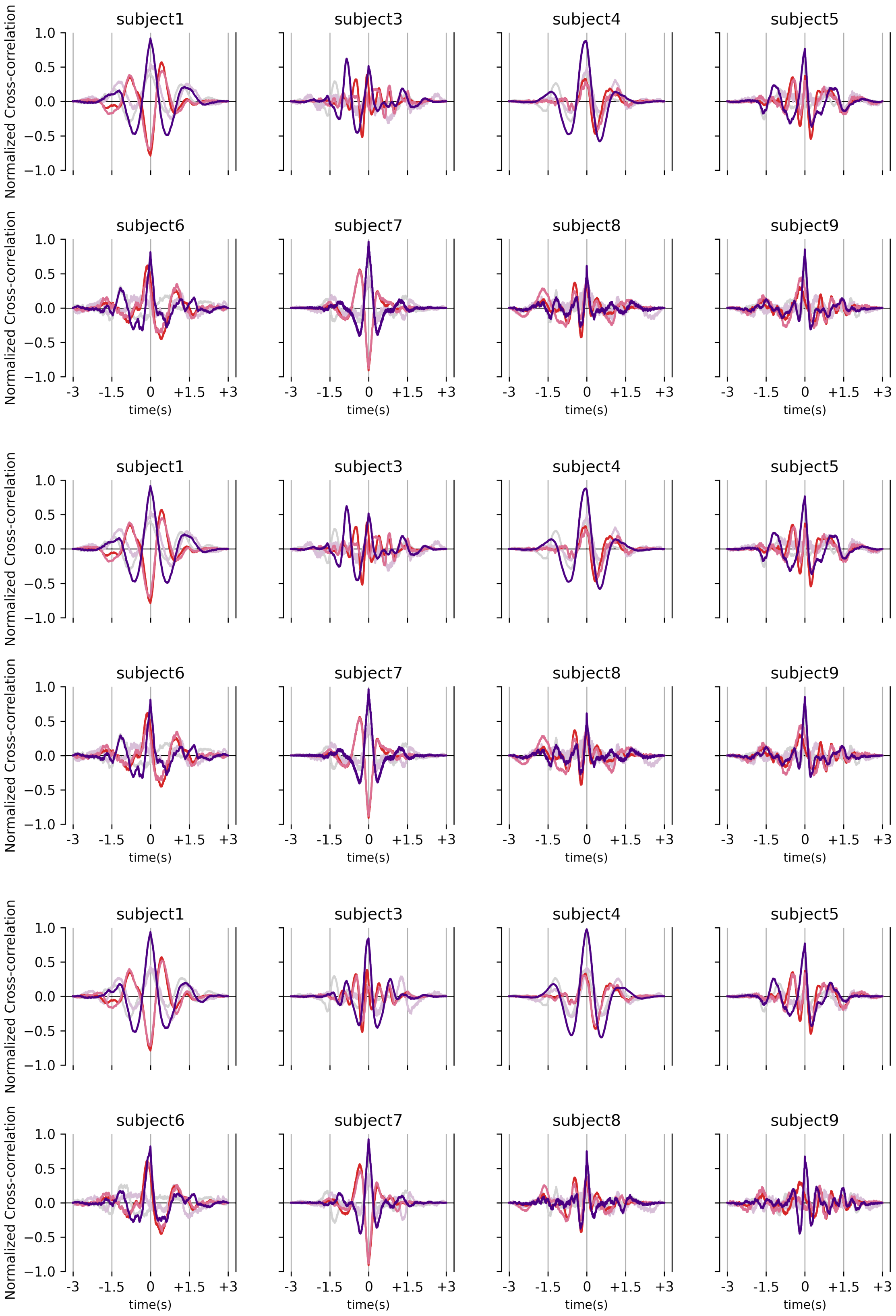


**Supplementary Figure 2.** The subject-wise full-time normalized cross-correlation between synthesized and actual brain signals in the arm-reaching dataset (Indigo:M1&Synth.M1, Red:Source&Synth.M1, Light gray: M1&Synth.M1(noise)). Top: mIPS, Middle: BA6d, and Bottom: MT

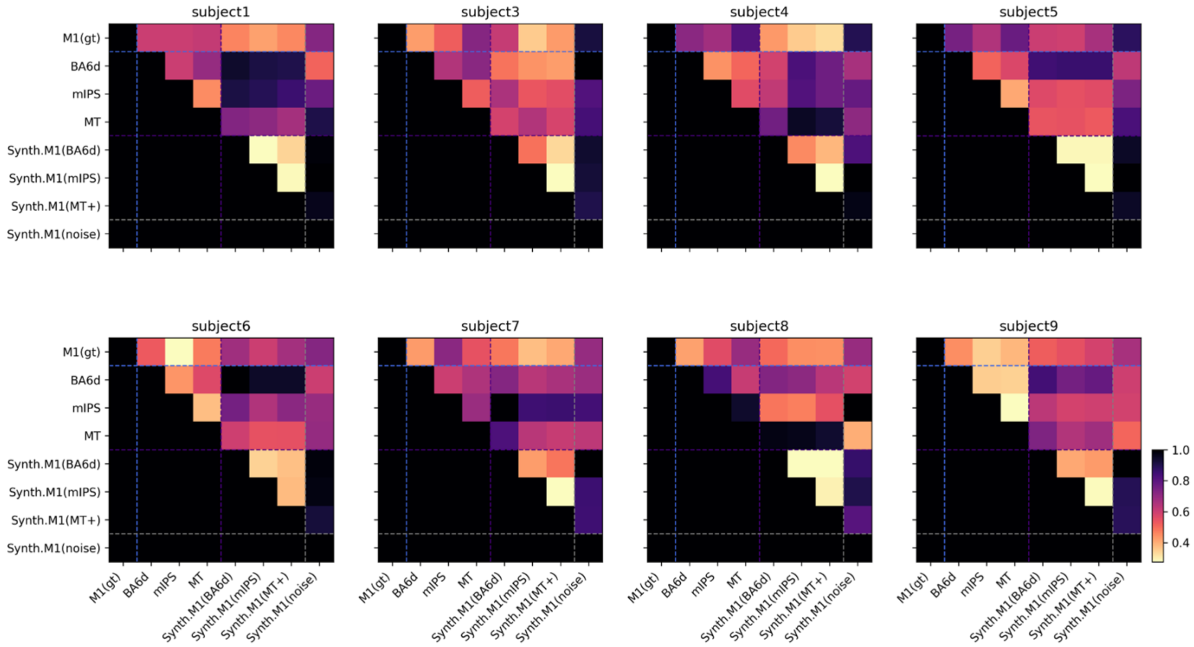


**Supplementary Figure 3.** The subject-wise normalized Fréchet distances (FDs) between the neural signal pairs.

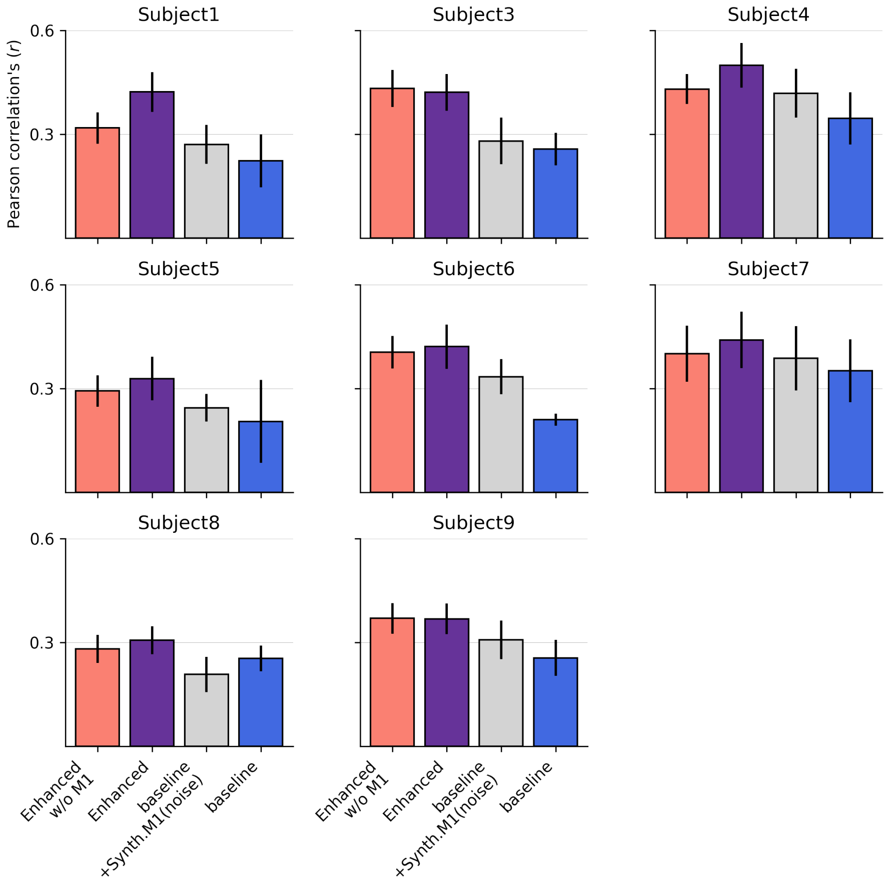


**Supplementary Figure 4.** The subject-wise arm-reaching movement decoding performance in each dataset condition
